# Immunogenicity and protective potency of Norovirus GII.17 virus-like particle-based vaccine

**DOI:** 10.1101/496190

**Authors:** Wei Chen, Tao Kang, Rongliang Yuan, Yuyang Zhang, Siqi Xin, Congwen Shao, Shenrong Jing

**Author notes:** Correspondence authors: Congwen Shao and Shenrong Jing. Address: Medical School, Kunming University of Science and Technology, No. 727, Southern Jingming Road, Chenggong District, Kunming, Yunnan, China 650500. Wei Chen and Tao Kang are equally contribution to the paper.

## Abstract

Noroviruses (NoVs) are a major cause of acute viral gastroenteritis in adults and children worldwide. Lacking of cell culture system and animals models that must be considered the virus like particles (VLPs) used as an effective vaccine development. In the present study, we investigated the expression of the major capsid protein (VP1) of Genogroup II, genotype 17 (GII.17) NoV using recombinant baculovirus system in insect cells and saliva binding blockade assay to detect their protective potency. Our results showed that GII.17 VLPs could be successfully generated in sf9 insect cells and electron microscopic revealed that GII.17 VLPs was visualized as spherical particles of −35nm in diameter. Immunized mouse with purified VLPs produced GII.17 specific sera and could efficiently block GII.17 VLPs binding to saliva histo-blood group antigens (HBGAs). Together, these results suggested that GII.17 VLPs represent a promising vaccine candidate against NoV GII.17 infection and strongly support further preclinical and clinical studies.

## Introduction

Human noroviruses (NoVs) infections are considered a significant public health problem, which cause more than 50% of non-bacterial gastroenteritis characterized by fever, vomiting and diarrhea emerging frequently, espically in adults and children [1, 2]. NoVs divided into 5 genogroups designated GI through to GV and subdivided into 9 and 22 genotypes which was first linked to human disease in 1972 [3, 4]. From the mid-1990s, Genogroup II, genotype 4 (GII.4) NoV have been predominantly in outbreaks and with every two or three years emergence of new variants [5]. However, since the emergence of the novel Genogroup II, genotype 17 (GII.17) Kawasaki_2014 strain, a new pandemic strain has predominant in the word [6]. The novel GII.17 not only caused NoVs outbreaks but also spread sporadically with cases reported in China (Shanghai, Taiwan) and USA during the winter of 2014_2015 [7–9]. From 2015, NoV GII.17 continued to circulate in many countries in South America, Europe and Asia [10–12]. These epidemiological surveys indicate that NoV GII.17 is becoming an important etiological agent of gastroenteritis.

NoVs belongs to the family of Caliciviridae and are nonenveloped, single-stranded, positive-sense ribonucleic acid (RNA) viruses. The genomes are 7.4-7.7 kb in size and have typically organized into three open reading frames (ORFs). The ORF1 encodes nonstructural proteins, such as the RNA dependent RNA polymerase (RdRp), while the structural proteins, such as the major and minor capsid protein (VP1 and VP2), are encoded by ORF2 and ORF3, respectively [13]. VP1 is a 60 KDa protein that can be divided into a shell domain (S domain) and a protruding domain (P domain). The S domain forms a scaffold surrounding the viral RNA and responsible for the shell structure of the capsid, whereas the P domain contains neutralizing epitopes and binds to histo-blood group antigens (HBGAs) receptors, responsible for the genetically variable, builds the viral spikes and facilitates cell attachment [14–16]. VP2, a minor capsid protein, exhibiting multiple functions, which is observed to stabilize and promote expression of VP1 [17].

The newly outbreak NoV GII.17 strain has been sweeping all over the world. However, no vaccines or specific treatments are available. Due to a lack of a permissive cell culture system and available an animal model, the main capsid protein VP1 formed the virus like particles (VLPs) has been assembled and used as NoVs vaccine candidates in preclinical and clinical studies [18]. VLPs have shown high immunogenicity, safety and promising results as human enteroviruses vaccines, such as enterovirus 71 (EV71), coxsackievirus A16 (CA16), and enterovirus D68 (EV-D68) [19–21].

In the present study, we constructed GII.17 VP1 and expressed using recombinant baculovirus expression system in sf9 cells which lead to formation of VLPs that are morphologically and antigenically similar to true virions. Our results showed that GII.17 VLPs could be readily produced in the baculovirus/insect cell system and these VLPs could induce potent neutralizing antibody responses and provided effective protection.

## Materials and Methods

### Construction of baculovirus expression vectors

The full-length capsid protein VP1 coding sequence (Genbank number, KT992785) was optimized based on insect cells expression and synthesized by Sangon Biotech (Shanghai, China) with flanked by *BamHI* and *Xba* l sites at its 5’and 3’ends, respectively. The optimized gene cloned into pFastBac-Dual-EGFP vector (our laboratory construction previous), named pFastBac-Dual-EGFP-VP1 (GII.17). Then the pFASTBac-Dual-EGFP-VP1 (GII.17) vector purified plasmid DNA transform into DH10Bac^™^ *E.coli* for transposition into the bacmid. At last, the correct recombinant bacmid DNA transfect into sf9 insect cells to produce recombinant baculovirus.

### Production and purification of NoV GII.17 VLPs

After transfection about 96h (the green fluorescence reaches the most), the budded virus released into the medium and collected the medium which was the P1 viral stock. Amplified baculoviral stock until P5 viral stock and collected cells.

Cells were washed and lysed in phosphate buffer solution (PBS) with ultrasound. The lysates supplemented with 0.5 mol/L NaCl and 10% (W/V) PEG 8000 mixed for 6h, and then centrifuged at 10,000 rpm for 30 min. The resultant precipitates were loaded onto a 20% sucrose cushion, and subjected to 10%-50% sucrose gradients for ultracentrifuged at 40,000 rpm for 6h. After ultracentrifuged, fractions were taken from top to bottom and then analyzed contained VLPs by SDS-PAGE. The uninfected sf9 cells were subjected to the same treatments as the negative control. The final purified proteins were dialysis with physiological saline at 4°C overnight and used for immunization experiments.

### Electron microscopy

Purified GII.17 VLPs was negatively stained with 0.5% aqueous uranyl acetate, and observed by transmission electron microscopy with Tecnai G2 Spirit at 120V (Thermo, USA).

### Mouse immunization

Mice were purchased from Kunming Medical University (Yunnan, China). All animal studies were approved by the Institutional Animal Care and Use Committee at the Kunming University of Science and Technology.

GII.17 VLPs or the negative control antigen were adsorbed to the aluminum hydroxide adjuvant with 1:1 by vortexing to a final volume 100 μL containing 50 μg of GII.17 VLPs for each injection. Groups of 6 female BALB/c mice (6-8weeks old) were intraperitoneally (i.p.) injected with antigen mixtures at weeks 0 and 4. Blood samples were collected from each immunized mouse after the final immunization, and putted at 37°C for 2h and 4°C for overnight.

### Serum antibody measurement assay

NoV GII.17 specific antibodies in immunized mouse sera were detected by indirect ELISA assay. 96-well plates were coated with 0.1 μg (100 μL)/well of GII.17 VLPs at 4°C for overnight, followed by blocking with 5% milk diluted in PBS containing 0.05% Tween-20 (PBS-T) at 37°C for 1h; then incubated at 37°C for 2 h with 100 μL/well of serum samples that individual serum samples were diluted in series in PBS-T. At last, incubation with goat anti-mouse IgG HPR (Abcam, Ab6789, UK) at 37° C for 1h. Between each step, the plates were washed three times with PBS-T. The absorbance was measured at 450 nm using a microplate reader (Thermo, USA).

### Western blot assay

Mouse sera collected were used to determine their reactivates on VP1 protein. In brief, the VLPs proteins separated by SDS-PAGE were transferred to PVDF membrane and detected using mouse anti-GII.17 VLPs serum at a dilution of 1:20000 in PBS-T and then HRP conjugated goat anti-rabbit IgG polyclonal antibody was added at 1:10000 in enzyme buffer. The membrane was developed after another three times wash using PBS-T with DAB.

### GII.17 VLPs-HBGAs binding assay in vitro

For the in vitro VLPs-HBGA binding assay, the eighty-four saliva samples collected from blood type A, B, AB and O individuals were used. The protocol for the GII.17 VLPs saliva HBGAs binding assay was conducted based on previously [22, 23]. Briefly, human saliva with known ABO antigens was boiled for 10 min and diluted at 1:1000 in PBS, added into 96-well plates (100 μL/well) and incubated at 4°C for overnight, followed by blocking with 5% milk diluted in PBS-T at 37°C for 1h. GII.17 VLP at 0.1 μg (100 μL)/well was incubated at 37°C for 2 h, followed serum samples that were diluted 1:1000 in PBS-T at 37°C for 2 h with 100 μL /well. At last, incubation with goat anti-mouse IgG HPR (Abcam, Ab6789, UK) at 37°C for 1h. Between each step, the plates were washed three times with PBS-T. The absorbance was measured at 450 nm using a microplate reader. At the same time, we used PBS-T without sera as a negative control.

### GII.17 VLPs-HBGAs binding blockade assay

The blocking activity of sera against GII.17 VLPs was determined by in vitro VLPs-HBGAs binding blockade assay. The protocol was performed basically the same as in vitro VLPs-HBGA binding assay except for serum dilution and antibody. Since GII.17 VLPs exhibited the strongest binding to blood type AB salivary HBGAs, a blood type A saliva sample was selected for the blockade assay. All reagents, sera dilutions and reaction times were optimized and conducted saliva-VLPs blocking assay using mouse sera. The 96-well microplates added to purified VLPs containing VLPs mouse serum at 37°C for 1 h, and rabbit anti-GII.17-P domain monoclonal antibodies (Taizhou SCIVAC Bio-Tech CO., Ltd) diluted was added. Wells added with VLPs containing without VLPs mouse serum was selected as control. The blocking index was calculated in % as (mean OD without VLP sera-mean OD with VLP sera)/mean OD without VLP sera × 100% [24].

### Statistical analysis of data

Statistical analyses were performed using Graphpad Prism v5 software. Antibody titers or OD values between groups were compared by Student’s two-tailed t-test.

## Results

### Expression and characterization of NoV GII.17 VLPs in sf9 cells

In order to confirm the expression of GII.17 VP1 protein in sf9 cells, EGFP and VP1 genes were together cloned into pFastBac-Dual vector under the control of pH and p10 promoters, respectively (Fig.1A). The recombinant baculovirus was using Bac-to-Bac baculovirus expression system and then infected insect cells. When the green fluorescence reaches the most under the fluorescence microscope, we collected cells. Sucrose gradient ultracentrifugation to purification the VP1 revealed by SDS-PAGE and the results which corresponds to the full-length VP1 protein of -58KDa (Fig.1B). The purity and integrity of VLPs were showed by transmission electron microscopy (Fig.1C). It should be noted that smaller, larger and full VLPs (about 35 nm) were observed.

**Fig. 1.**
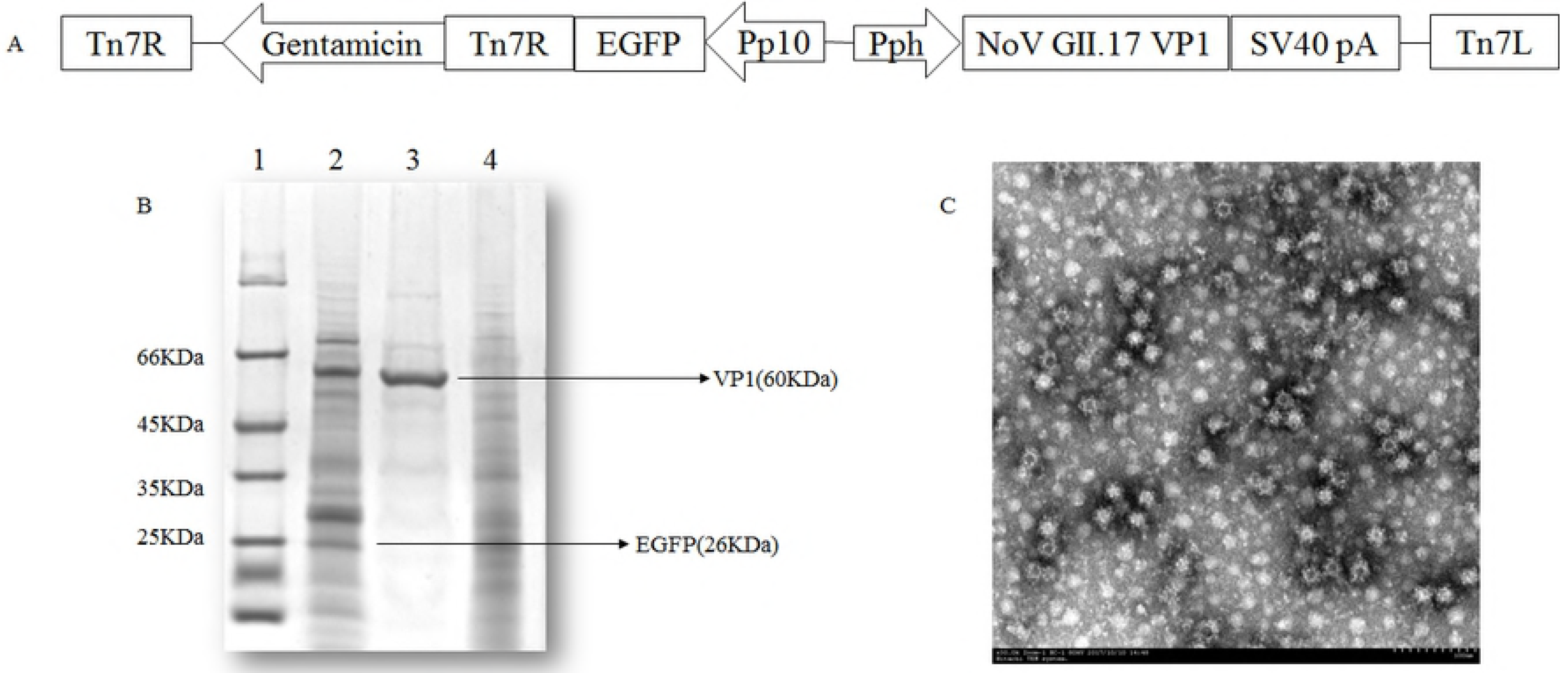
Co-expression of NoV GII.17 VP1 and EGFP in insert cell

### Specificity of the antibody response following immunization

To determine the immunogenicity, GII.17 VLPs was used to i.p. immunize BALB/c mice three at four weeks. Another group of mice was injected with PBS as a control. Serum samples were collected from two weeks after the final immunization and subjected to ELISA analysis for antibody measurement. All sera from VLPs immunized mice exhibited high binding activities towards GII.17 VLPs, whereas the control group did not exhibited significant reactivity (Fig. 2A). As showed in Western blot analysis (Fig. 2B) can confirmed the expression of VP1 protein. These results indicate that immunization with the VLPs but not the control antigen could induce GII.17 VLPs specific antibody responses in mice.

**Fig. 2.**
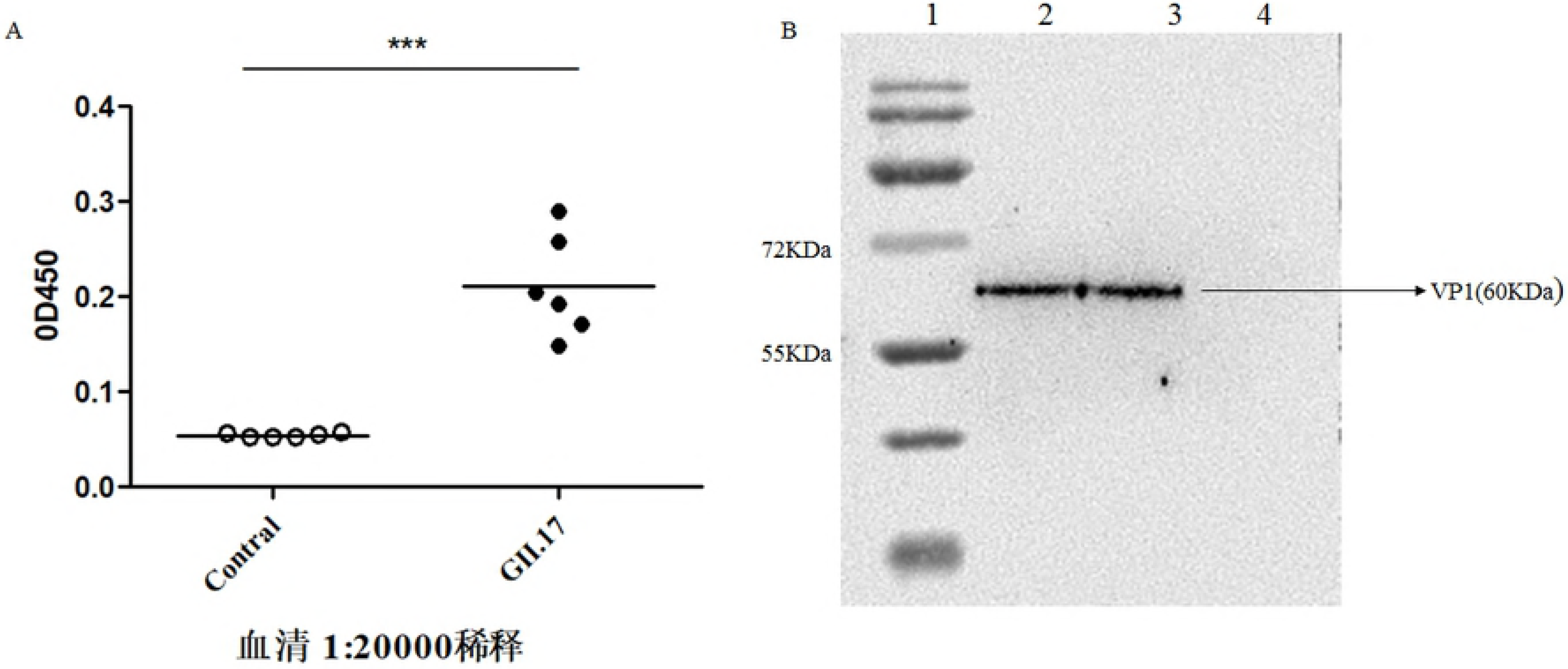
Serum antibody response in mice after immunization with GII.17 VLPs

### Assembled VLPs bind to salivary HBGAs

An in vitro VLP-HBGA binding assay was used to characterize the binding profiles of GII.17 capsid protein assembled VLPs. As shown in Fig. 3, VLPs bound to blood type A, B and AB salivary HBGAs, respectively, while weak binding to type O salivary HBGA.

**Fig. 3.**
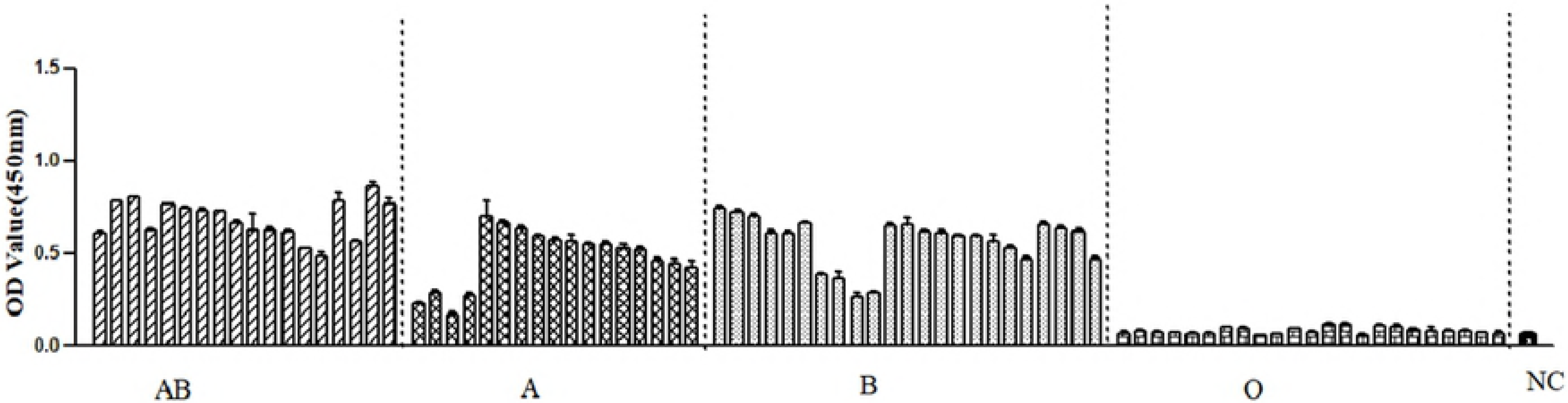
Characterization of binding profile of GII.17 VLPs to blood type A, B and AB salivary HBGAs, respectivly

### The capacity of VLPs immunized mouse sera to block GII.17 VLPs saliva binding

As we known, the protective function of anti-NoV serum antibodies correlates with their capacity to block NoV VLPs binding to HBGAs, therefore, the serum antibody-blocking assay was conducted to test the NoV neutralization [25–27]. We collected the saliva HBGAs, including A, B and O antigens and has therefore been used in the blockade assay. We used rabbit sera against GII.17 P domain as antibody in our study. Saliva sample collected from a blood type AB individual was used in the in vitro VLPs-HBGA binding blockade assay which represented the highest binding signal. As was shown in Fig. 4, the blocking indices were higher for antisera from blood type AB. Addition of serum against GII.17 VLPs blocked the binding of GII.17-VP1 capsid protein assembled VLPs to salivary HBGAs.

**Fig. 4.**
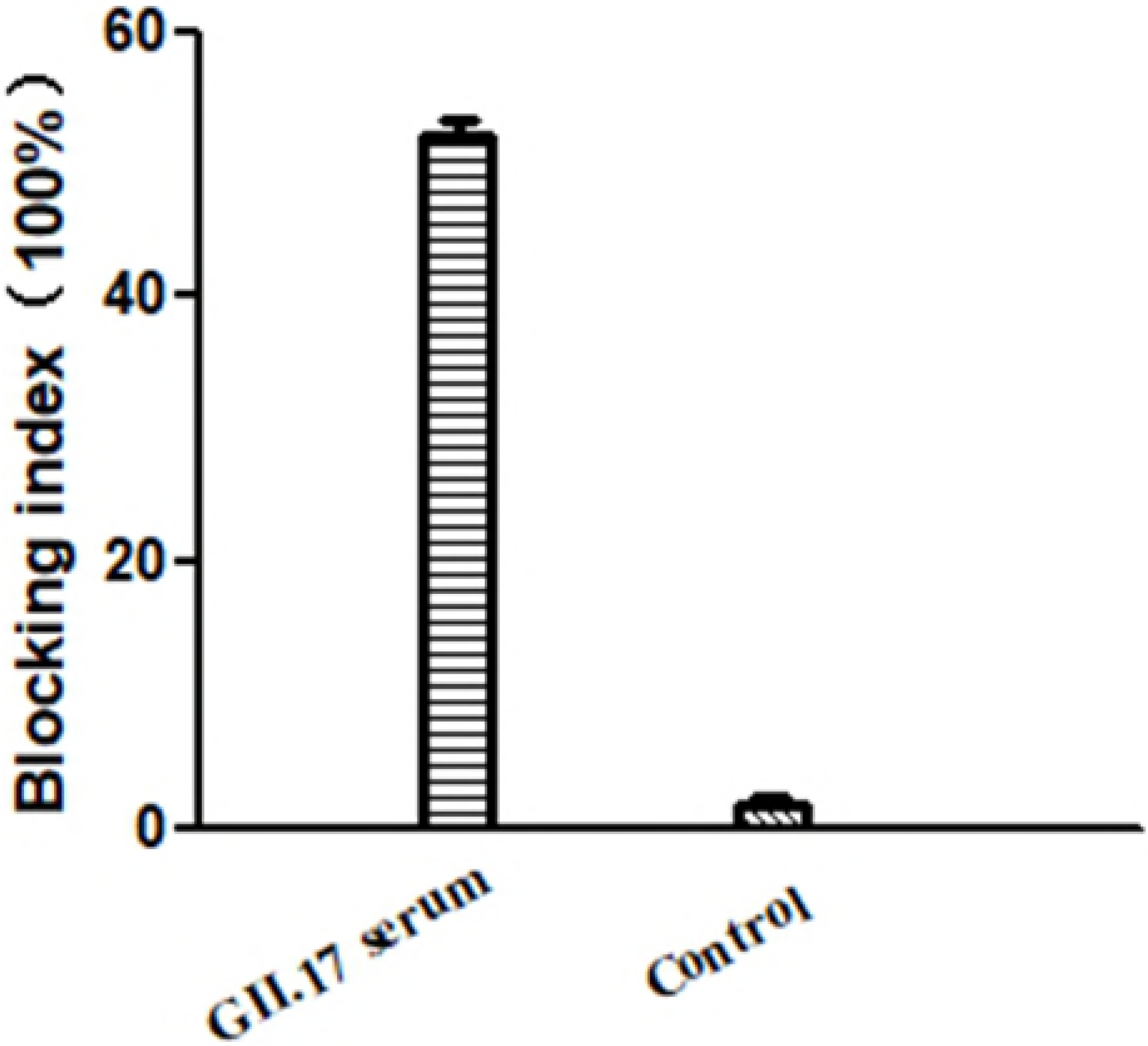
VLP-HBGAs binding blockade assay

## Discussion

NoVs are the major cause of acute non-bacterial gastroenteritis all over the world especially in children. As rapid evolution of NoV GII.4 with new strain replacing previous strain every two or three years, the newly outbreak NoV GII.17 strain has been sweeping all over the world. Thus, it is highly desirable to formulate an effective and rapid NoV GII.17 vaccine to achieve. In our study, we represented that the fist insect cell expressed GII.17 VLPs is promising vaccine.

The NoVs capsid protein is divided into two domains: S domain, forming the core of the virus-like particle, and the P domain, mainly involved in receptor binding [14]. Comparison of avidity of IgG from mice immunized from VLPs and P particles, the results suggested that significant higher avidity for VLPs immunized mice and also showed immunization route and adjuvants have no effect on the IgG avidity [28]. Some of NoV VLPs candidates have shown efficacy in human trials [26, 29]. As we known, the VP1 which can self-assemble into VLPs when expressed in sf9 cells using recombinant baculovirus expression system [30]. Therefore, in our research, GII.17 VLPs has been regarded as the most promising vaccine candidates and were intraperitoneally (i.p.) injected with antigen mixture with aluminum hydroxide adjuvant.

Due to the lack of an animal model available for human NoVs infection, challenge study cannot be conducted to assess the efficacy of immunization. In recently, B cells as a cellular target of human NoVs and could be development of an in vitro infection model [31]. Prior to that, as we known, the HBGAs as important receptors for NoVs infection and the HBGAs-VLPs blocking assay has been used as a surrogate method for evaluation of NoVs neutralization in vitro.

Among the GII.17 VP1, the P region is further divided into the P1 and the P2 subregion. The P2 subregion is located at the outermost layer of the entire capsid, which is important for the binding of NoVs and antibody recognition [32]. In *Escherichia coli,* expression of P domain can lead to formation of P-particle, a 12- or 24 dimers, which contained all the elements required for receptor binding and had been possessed the immunogenic in animal studies [33, 34]. So to characterize the binding specificity of GII.17 VP1 capsid protein assembled VLPs, we used rabbit sera against GII.17 P domain in our study.

HBGAs have been proposed as receptors for NoVs. Due to the NoVs different genotypes, such as GI.3 and GII.2 did not bind to synthetic or salivary HBGAs, whereas GII.4 could bind to salivary HBGAs [35, 36]. In our study, VLPs from capsid proteins of epidemic NoV GII.17 exhibit broader HBGAs recognition except blood type O individual, but lacking of the reasons to explain. Maybe the same as another study reported that the recombinant MxV (GII.3) VLPs demonstrated binding to salivary HBGAs from a blood type A individual [37].

At the same time, we used human saliva to evaluate in the blockade assay. Our results indicate that rabbit serum against GII.17 VLPs blocked the binding of capsid protein assembled VLPs to salivary HBGAs from a blood type AB individual and blocking rate up to 60%, whereas the anti-PBS did not have such effects. The discovery of new receptors or factors that promote the binding of GII.17 VLPs to HBGAs is a key for following vaccine development and vaccine efficacy evaluation. These data also suggest important information about the presence of other factors involved in the binding of GII.17 VLPs to HBGAs.

## Conclusion

Considering the facts of prevalence GII.17, the NoVs vaccine should be urgently developed. Our results provided that the GII.17 VLPs induced antibodies should be protective against in vivo NoV GII.17 infection. Therefore, we identified GII.17 VLPs as a viable candidate. NoVs vaccine represents a step forward in developing broad-spectrum, multivalent vaccines against acute viral gastroenteritis.

## Acknowledgments

This research is supported by grants from the National Natural Science Foundation of China (No. 81860357) and Yunnan Provincial Department of Education (2017ZZX138).

## Author Contributions

Conceptualization: Congwen Shao, Shenrong Jing

Data curation: Rongliang Yuan, Yuyang Zhang, Siqi Xin

Methodology: Wei Chen, Tao Kang, Yuyang Zhang

Project administration: Congwen Shao, Shenrong Jing

Supervision: Shenrong Jing

Writing-original draft: Wei chen

Writing-review&editing: Shenrong Jing

